# Development and Optimisation of a Defined High Cell Density Yeast Medium

**DOI:** 10.1101/846006

**Authors:** Tania Michelle Roberts, Hans-Michael Kaltenbach, Fabian Rudolf

## Abstract

*Saccharomyces cerevisiae* cells grown in a small volume of a defined media neither reach the desired cell density nor grow at a fast enough rate to scale down the volume and increase the sample number of classical biochemical assays, as the detection limit of the readout often requires a high number of cells as an input. To ameliorate this problem, we developed and optimised a new high cell density (HCD) medium for *S. cerevisiae*. Starting from a widely-used synthetic medium composition, we systematically varied the concentrations of all components without the addition of other compounds. We used response surface methodology (RSM) to develop and optimise the five components of the medium: glucose, yeast nitrogen base, amino acids, mono-sodium glutamate and inositol. We monitored growth, cell number and cell size to ensure that the optimisation was towards a greater density of cells rather than just towards an increase in biomass (i.e larger cells). Cells grown in the final medium, HCD, exhibit growth more similar to the complex medium YPD than to the synthetic medium SD, while the final cell density prior to the diauxic shift is increased about three- and tenfold, respectively. We found normal cell-cycle behaviour throughout the growth phases by monitoring DNA content and protein expression using fluorescent reporters. We also ensured that HCD media could be used with a variety of strains and that they allow selection for all common yeast auxotrophic markers.

## Introduction

The yeast *Saccharomyces cerevisiae* is a broadly used eukaryotic model organism in basic research as well as an important tool in biotechnology (Botstein and Fink, 2011). In contrast to many other model organisms, yeasts offer comparatively simple genetic manipulation using either plasmid-based expression or genome modifications (Gietz and Woods, 2016). Over the last 70 years, the community established a series of commonly used selection markers for this purpose, but their use requires a defined growth medium for auxotrophic selection (Gnügge and Rudolf, 2017). Most commonly, a defined growth medium uses glucose as its only carbon source with concentrations varying between 10 g/*ℓ* and 60 g/*ℓ*. Additionally, a nitrogen source —in most cases ammonium sulphate— phosphate and a combination of vitamins are required. The latter components all come from yeast nitrogen base (YNB), which might need to be supplemented with inositol for efficient growth of certain lab strains (Hanscho et al., 2012). Lastly, the most common defined media have a set of amino acids added, whose composition and concentrations vary among recipes and laboratories (Hanscho et al., 2012). While these media have a clearly defined composition, the cell density at the diauxic shift is lower and the growth rate of *S. cerevisiae* is slower compared to cells grown in the complex medium YPD. The growth rate varies during the pre-diauxic shift growth and, therefore, balanced growth is only obtained in a short window.

The majority of work with *S. cerevisiae* is performed on agar plates when scoring colony growth, or in shaking cultures for single cell and biochemical assays. While single cell assays only require a small culture volume, biochemical assays require starting culture volumes ranging between 5 and 500 m*ℓ* of medium. Increasing the throughput of agar work and single cell assays can be easily achieved using arrayed colonies or liquid culture on standard SBS format plates (Costanzo et al., 2016). Scale-up of the throughput of liquid experiments, however, depends heavily on the sensitivity of the readout and ease of cell lysis (Ovalle et al., 1999). Prime examples for successful miniaturisation are transcriptional reporter assays where a readout of interest can be induced and read by sensitive methods such as *β*-galactosidase, or scoring of single cell fluorescence using flow cytometry (Rajasarkka and Virta, 2012; Petrovic et al., 2017). Meanwhile, miniaturisation of classical biochemical assays is more daunting. For example, 1.7 m*ℓ* of culture needed to be grown in the complex medium YPD for a simple study of the level of expressed protein using western blotting with the sensitive TAP tag (Ghaemmaghami et al., 2003). Even larger volumes are required for co-immunoprecipitation using the TAP tag or any other tag (Suter et al., 2007), except when the analysis is performed using sensitive, state-of-the-art mass spectrometry (Gavin et al., 2002, 2006). For the vast majority of labs, miniaturisation and small-scale scale up of assays such as western blotting, co-immunoprecipitation or enzyme activity measurements are therefore unattainable, especially when proteins are expressed at low levels.

A crucial step towards miniaturisation of all biochemical assays is the increase in the amount of starting material by increasing the cell density at the diauxic shift, without dramatic changes to the cell physiology. Preferably, this should be achieved by optimisation of the concentrations of the ingredients of commonly used defined growth medium, while keeping the growth rate, cell size and cell cycle timing similar to the established standard conditions to not dramatically alter the cellular physiology. Such a medium would aid to lift the readout above the detection limit and thereby allow more labs to perform biochemical assays using standard SBS-plates.

Optimising growth conditions is a standard procedure to make biotechnological production processes economically viable, mainly as more biomass per volume requires less production capacity, generates less waste, and gives more yield per volume (Westman and Franzén, 2015). However, most of these optimisation start from a complex medium and then add buffer substances, carbon sources and vitamins to increase biomass (Pereira et al., 2013; Wang et al., 2010; Mendes-Ferreira et al., 2010). The resulting media change the cellular physiology and do not allow for the use of auxotrophic markers and are therefore ill-suited for basic research. Only few media were optimised for use with auxotrophic markers and allow balanced growth during the exponential phase. An interesting example of this type increases secretion of proteins, but does not increase the cell density by adding additional amino acids to the classical defined synthetic medium while keeping the rest of the ingredients constant (Wittrup and Benig, 1994; Görgens et al., 2005).

*response surface methodology (RSM)* is a statistical technique for iterative improvement of a process by sequential experimentation and is a common technique for optimising (bio-)chemical and industrial processes (Box and Hunter, 1957; Box et al., 2005). The response surface describes the expected response of the process (such as the cell density) for any specific values of a set of explanatory variables (such as the concentration of components of a medium). The goal of RSM is to find a (local) optimum, that is, a combination of concentrations of components that maximise the response. In general, the response surface is unknown and RSM iterates between exploration around a defined point to locally approximate the surface and measure the response along the predicted path of steepest ascent. In a first set of experiments, the response surface is explored in the vicinity of an initial medium composition (Figure 1A). For this, a set of new compositions is designed by systematically and simultaneously altering the amounts of all components, and the resulting responses are measured. A regression model is then fitted to the measurements to locally approximate the response surface and to calculate the direction that yields the steepest ascent on this surface (the *canonical path*). In a subsequent set of experiments, the responses are measured along this path at different distances from the initial medium (Figure 1B). Typically, the condition with highest response is then taken as the new intermediate medium composition and the two steps are repeated (Figure 1B). This procedure is continued until the resulting response is deemed optimal, e.g. no increase in the response is gained, the resulting response is deemed sufficiently high, or the experimental resources are exhausted.

**Figure 1.**
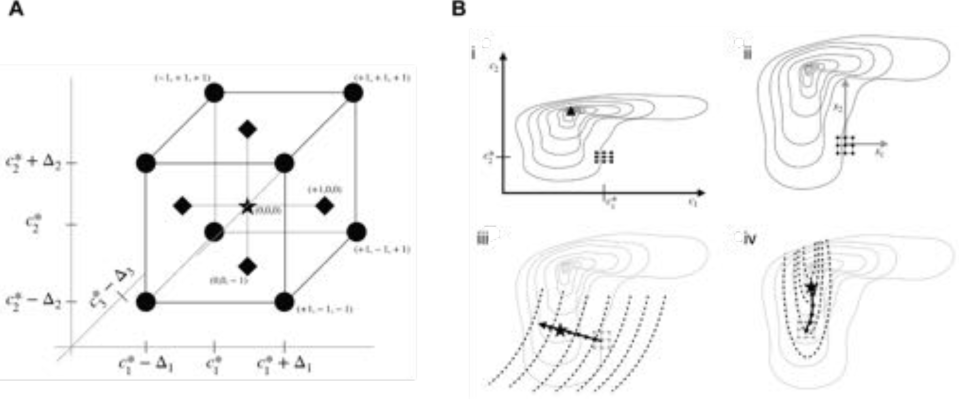
Overview of RSM methodology. (A) Central composite design for three components. Star: centre point, circles: factorial points, diamonds: axial points. For clarity, axial points for third component are not shown. After coordinate transformation, centre point is at origin and factorial points are (±1, ±1, ±1); new coordinates are shown for select points. (B) Sequential optimisation using response surface method. i: Original medium components with concentrations *c*_1_ and *c*_2_, respectively, lead to response surface shown as thin solid lines with a maximum response denoted by a triangular point. The starting composition is 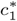 and 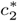, shown as a black point. ii: In the first exploration, RSM shifts and scales the coordinate system to *x*_1_ and *x*_2_ (grey lines). The central composite design requires measuring the response at the indicated round points. iii: dashed lines show the approximate response surface based on the locally fitted model. The CCD and the true surface are shown in grey. Following the canonical path (solid line) yields the intermediate optimum at the star-shaped point. iv: the procedure is repeated from the new intermediate optimum. The new local approximation predicts a maximum upward and to the left of the actual optimum, but the slightly curved canonical path still approaches the actual maximum. The procedure can be repeated again if required.

Here we present an optimised synthetic well-defined medium allowing for growth of *S. cerevisiae* lab strains to high cell densities. Only the composition of the commonly used synthetic SD medium is changed and it is therefore usable with common auxotrophic or drug resistance markers. We term it *high cell density (HCD)* medium. It yields tenfold more cells at the diauxic shift compared to SD medium and threefold compared to YPD. Importantly, it shows balanced growth during 12h of cultivation as scored by growth rate and the cell-to-cell variability of growth rate sensitive reporter genes. The medium was obtained by using two iterations of RSM-based designs. We only optimise the concentrations of the ingredients of the commonly used synthetic minimal media, with monosodium glutamate as a nitrogen source to allow for use of drug resistance marker genes. The RSM-based optimisation lead to non-intuitive changes to the medium composition: the medium was not only optimised for more glucose, but the amount of amino acids and YNB were also dramatically increased, while the nitrogen source, monosodium glutamate, was only slightly increased. We confirmed that the media are broadly applicable, using a series of auxotrophic markers as well as lab strains. HCD media can be used to amend and miniaturise biochemical assays, such that they can be performed in 96-well deep-well plates. Additionally, the media are favourable for protein production and especially protein secretion, as shown by an increased and constant per-cell expression of the test protein amylase during the whole growth phase.

## Results

### Optimisation using experimental design

In order to apply RSM, we need to specify an initial medium composition and the maximal deviations from this composition for exploring the response surface. We chose our initial medium based on a YNB supplemented version of a previously published optimised yeast medium to optimally support growth of the BY strain series (Hanscho et al., 2012). We reasoned that this medium provides the lowest growth level and therefore chose the starting conditions with the published medium at the lowest considered concentrations. Therefore, our first exploration only incorporated increases in concentrations.

We decided to optimise growth of the widely-used BY4741 strain background, using a mata strain, bearing *his3*Δ, *leu2*Δ, *ura3*Δ, and *met15*Δ gene deletions. We verified the suitability of our starting medium and the reproducibility of the assay by examining growth of cells of six independently generated cultures (Figure 2A). This simple growth test revealed that our starting medium already outperforms the standard medium and that our readout is sufficiently reproducible for performing a response surface optimisation.

**Figure 2.**
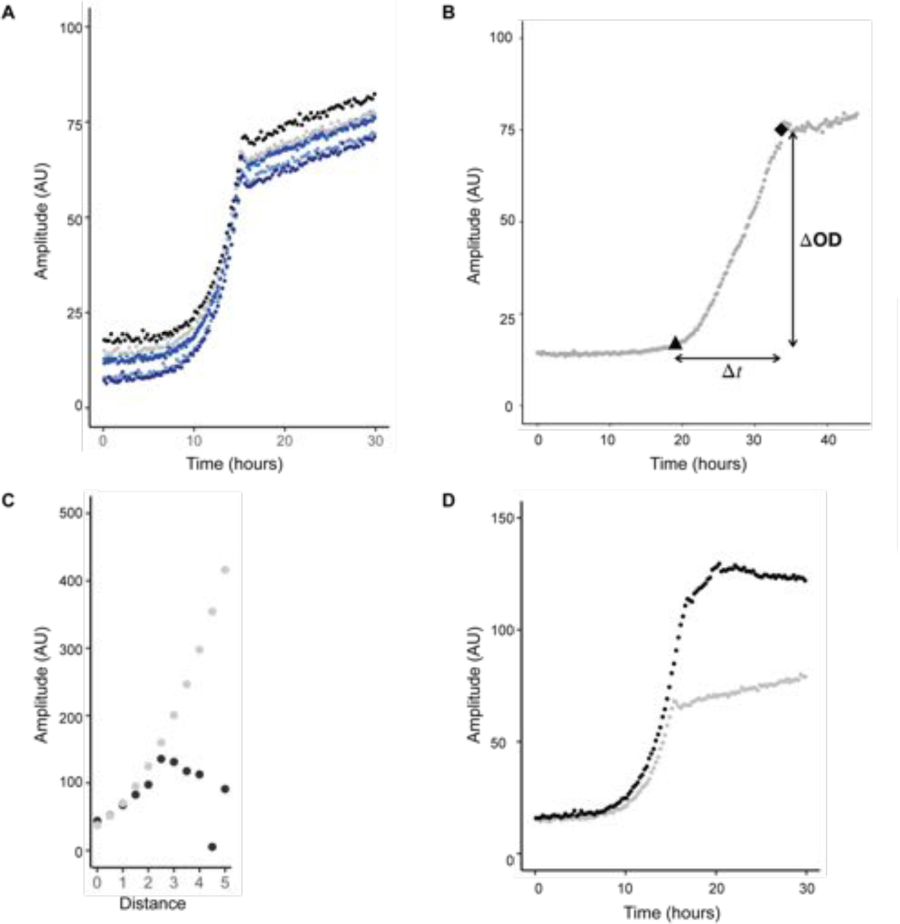
Utilisation of RSM to optimise media for increase in OD. (A) A defined yeast minimal media was used as the starting condition and centre point for RSM, 6 replicates are shown. (B) For the first iteration the increase (ΔOD) in optical density (Amplitude) from beginning of growth (triangle) to onset of the diauxic shift (diamond) was optimised. (C) Comparison of predicted (grey) and measured (black) increase in optical density (Amplitude) during growth along the first gradient of steepest ascent. (D) Comparison of growth of BY4741 cells in the starting conditions (grey) and the optimal conditions (black) after the first round of optimisation.

Next, we needed to simplify the experimental design, as fifteen components would lead to an infeasibly large experiment. We created five groups of components: (i) glucose concentration (C-source), (ii) monosodium glutamate as nitrogen source to keep the medium compatible with drug resistance markers, (iii) a mix of amino acids supplementing growth together with additional amino acids required for fast growth of BY strains, (iv) YNB without amino acids or ammonium sulphate, and (v) inositol. The initial amounts for each component and the low and high values considered for exploration are shown in Table 1 leading to a design with 32 experimental conditions; the initial optimised medium corresponds to the condition in the row labelled “Starting”.

**Table 1.** Media composition throughout the optimisation. First three rows: initial medium composition and the low and high points for the response surface method. Next row: intermediate optimum after first and second iteration. Last row: final optimum.

| Medium | Glucose | MSG | AA pool | YNB | Inositol |
| --- | --- | --- | --- | --- | --- |
| Starting | 4 % | 2 mg/ml | 1× | 3 mg/ml | 1 mg/ml |
| Low | 2 % | 1 mg/ml | 0.5× | 2 mg/ml | 0 mg/ml |
| High | 6 % | 3 mg/ml | 2× | 4 mg/ml | 2 mg/ml |
| Intermediate | 3 % | 2.7 mg/ml | 3.3× | 2.7 mg/ml | 1.4 mg/ml |
| Final | 6 % | 4.5 mg/ml | 6.0× | 3 mg/ml | 1.8 mg/ml |

To explore the response surface around our starting medium, we measured cell growth for each of the 32 design points using a Biolector growth reader (Figure S1). We used the increase in amplitude (or optical density) ΔOD (Figure 2B) from the start of growth to the beginning of the diauxic shift as the response that we aimed to maximise. This gives a crude but sufficient approximation of the increase in cell number. Six out of 26 conditions indeed showed an increase in biomass, while seven showed a discernible decrease. The remaining 13 showed similar growth as the starting medium.

After performing the exploration (Figure S1), we calculated the path of steepest ascent and measured the response along it (Table S1). We found that the predicted growth versus the measured growth matched well for the first six points along the canonical path (Figure 2C), after which the approximation of the response surface breaks down and the measured growth starts to decrease. Therefore, we took the sixth point as our intermediate optimum; it corresponds to the composition shown in row “Intermediate” of Table 1. A comparison of the growth curves between the starting medium and this optimum of this first round of RSM is shown in (Figure 2D). Interestingly, “Intermediate” showed a threefold increase in the amino acid pool, a slight increase in MSG and a decrease in glucose and YNB compared to our starting condition.

In order to ensure that the direction of the optimisation was indeed towards more cells and not simply towards more biomass, we checked if the cells behaved normally with regards to cell size, and measured cell size at the end of growth in the Biolector using a Coulter counter. We found no indication of abnormal cell sizes up to the “Intermediate” condition and a slight increase thereafter. We also did not find any obvious correlation between cell size profile and a specific medium component, as was reported for inositol (Hanscho et al., 2012), suggesting that it is a combination of several components that affect cell size distribution.

Using the “Intermediate” composition as our new starting point, we iterated the two steps of response surface exploration and following the canonical path again and obtained a second, increased response shown in Table S2. To ensure reproducibility of our results, we additionally performed two independent replicates of the exploration and subsequent experiments to pursue the direction of steepest ascent. Both replicates are in excellent agreement, as are the predicted and experimental responses (Figure 3A). The vertical shift between predictions and measurements is likely due to differences in calibration of the growth reader between the first and second round of optimisation, which we performed several weeks apart.

**Figure 3.**
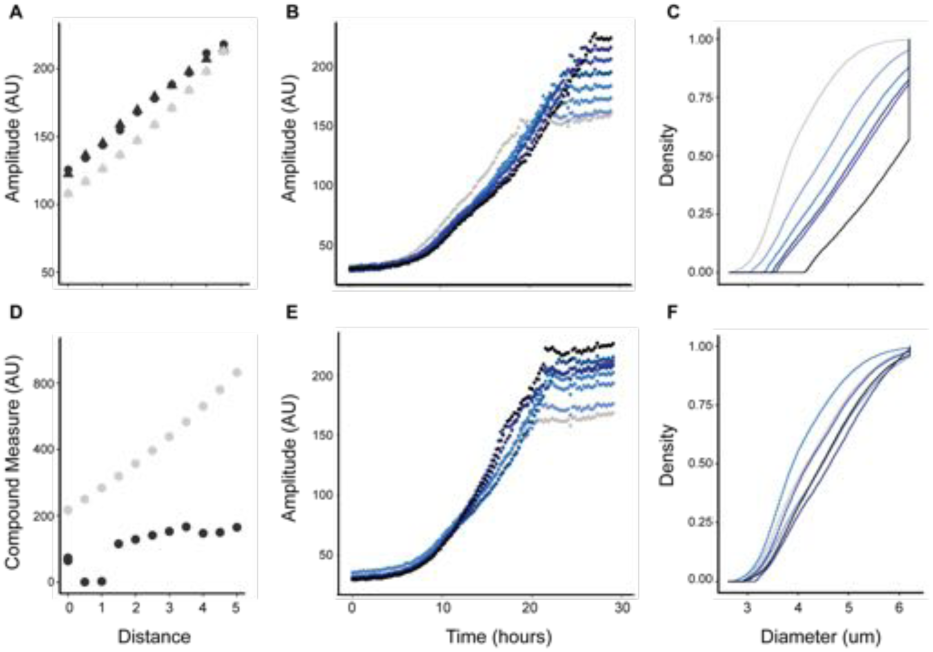
Comparison of gradient pursuit for optical density (OD) only or OD and time. (A) Comparison of predicted (grey) and measured (black) increase in OD (Amplitude) during growth along the second gradient of steepest ascent. The two point shapes correspond to two independent replicates of exploration and gradient pursuit experiments. (D) Comparison of predicted (grey) and measured (black) responses of increase in optical density adjusted for duration (“compound measure”) along modified gradient. The corresponding growth curves of BY4741 cells are shown in panel B and E, respectively. (C, F) The cell size of BY4741 cells from logarithmically growing cultures was measured using a Coulter counter. The empirical cumulative distribution function of the measured cell size of the optimisation based on OD (C) or OD and time (F).

In contrast to the first iteration, however, we found that higher responses were associated with slower growth and increased cell size (Figure 3B, C), suggesting that this second iteration of the optimisation was focused towards increasing total biomass rather than total cell number. In order to see if we could optimise for higher cell numbers, we repeated the calculation of the canonical path from the data of the second exploration. We used an adapted response function *y* =ΔOD − 2 Δ*t* to include the time Δ*t* elapsed between initiation of growth and reaching the diauxic shift as a crude measure for growth rate (Figure 2B). This new response function favours large increases in optical density, but additionally penalises long growth intervals. We verified that this steepest ascent prediction indeed follows a desired path by checking that no conditions in the vicinity of the path showed a decreased maximum growth, growth rate, or increased cell size.

We then measured the response of the ten predictions shown in Table S3 using the Biolector and found a clear increase in growth (Figure 3D). Up until a distance of 3.0 from the centre point, maximal biomass increased while cell size and growth rate stayed the same (Figure 3E, F). From distance 3.5 on, the compound measure, the cell size or both were less favourable than at distance 3.0. Similar to the initial second path, we observed that the predicted responses along the new path were overly optimistic. Nevertheless, the measured responses followed a clear upward trend, indicating that the corresponding medium composition lead to higher cell numbers. The medium at distance 3.0 approximately doubled the amount of glucose, MSG, amino acid pool and inositol, while only slightly increasing YNB compared to the starting point “Intermediate”.

We stopped the optimisation after this second iteration with modified response, and slightly modified the composition toward distance 2.5 to make it easier to pipette at the bench. We named the new medium *HCD* for *high-cell density* medium. Its composition is shown in Table 2 and its stock solution in Table 4.

**Table 2.** HCD Medium Composition. The crucial components are listed in table. Additionally, the medium has 20mM MES buffer pH 6 and traces of sulfate and phosphate. The detailed receipe is in Table 4

| Component | Final Concentration |
| --- | --- |
| Glucose | 6 % |
| MSG | 4.5 mg/ml |
| YNB | 12 mg/ml |
| Inositol | 1.8 mg/ml |
| **AA pool** |  |
| His | 210 µg/ml |
| Leu | 660 µg/ml |
| Lys | 720 µg/ml |
| Met | 240 µg/ml |
| Phe | 300 µg/ml |
| Ser | 225 µg/ml |
| Thr | 120 µg/ml |
| Ura | 240 µg/ml |

**Table 3.**
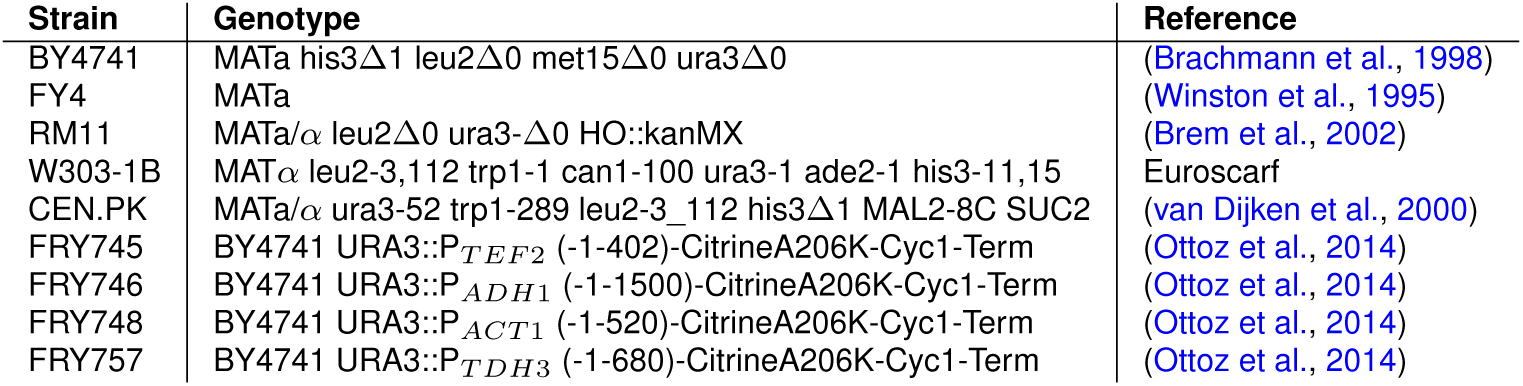
Yeast Strains

**Table 4.**
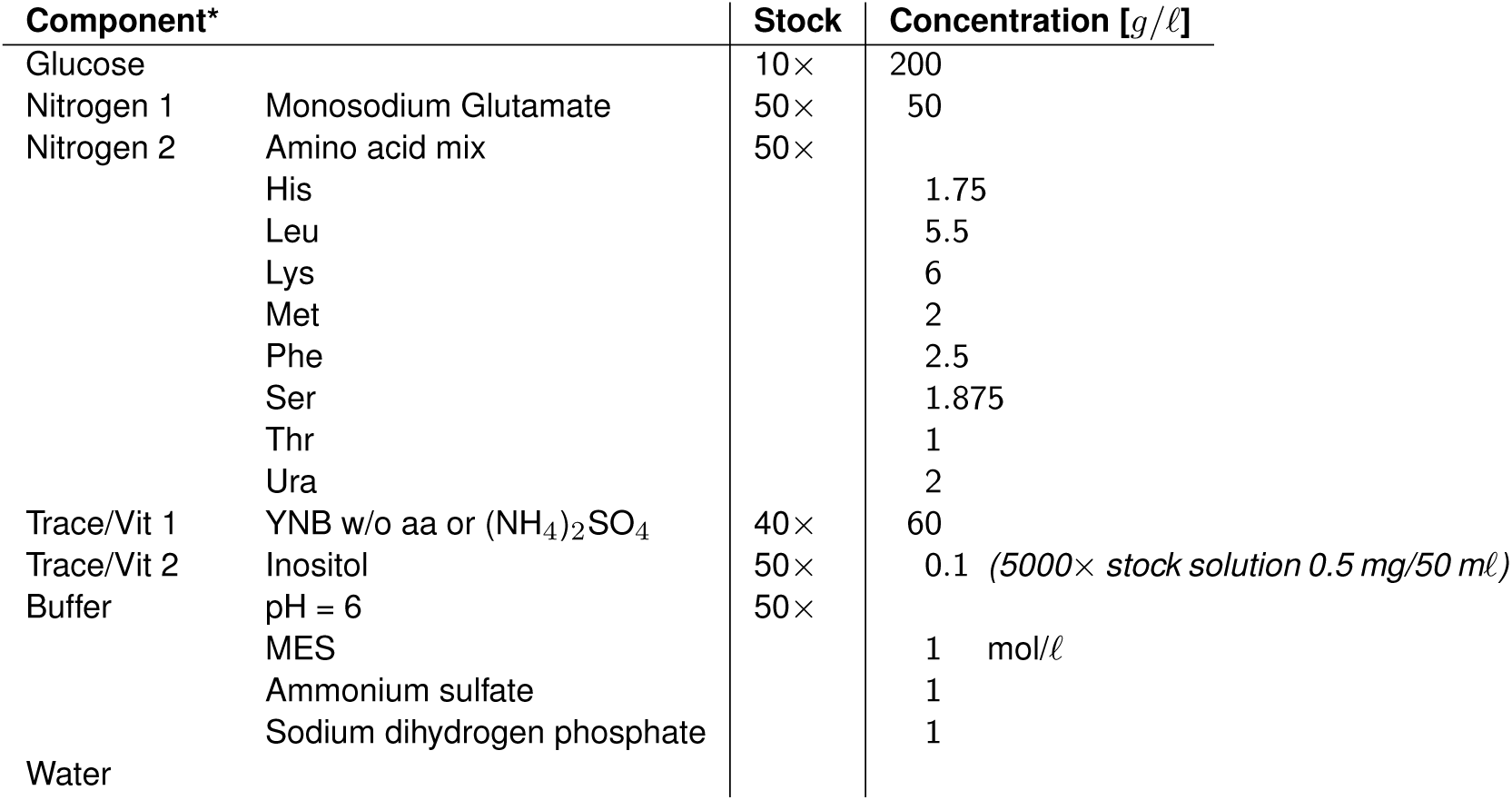
Stock solution composition used throughout the optimisation

### Cells grown in HCD medium exhibit normal physiology

HCD medium achieves cell numbers of 1.7 × 10^8^ cells/m*ℓ* at the end of the exponential phase of growth, a tenfold increase compared to the 1.6 × 10^7^ cells/m*ℓ* for SD, and a threefold increase compared to the 6.0 × 10^7^ cells/m*ℓ* for YPD. A direct comparison of typical growth curves and the cell size distribution for the three media is shown in Figures 4A, B, S2. It shows similar growth rates of HCD and YPD, while extending the balanced growth phase from 5 to 12h and that the cumulative cell size distribution of strains grown in three different media are almost identical.

**Figure 4.**
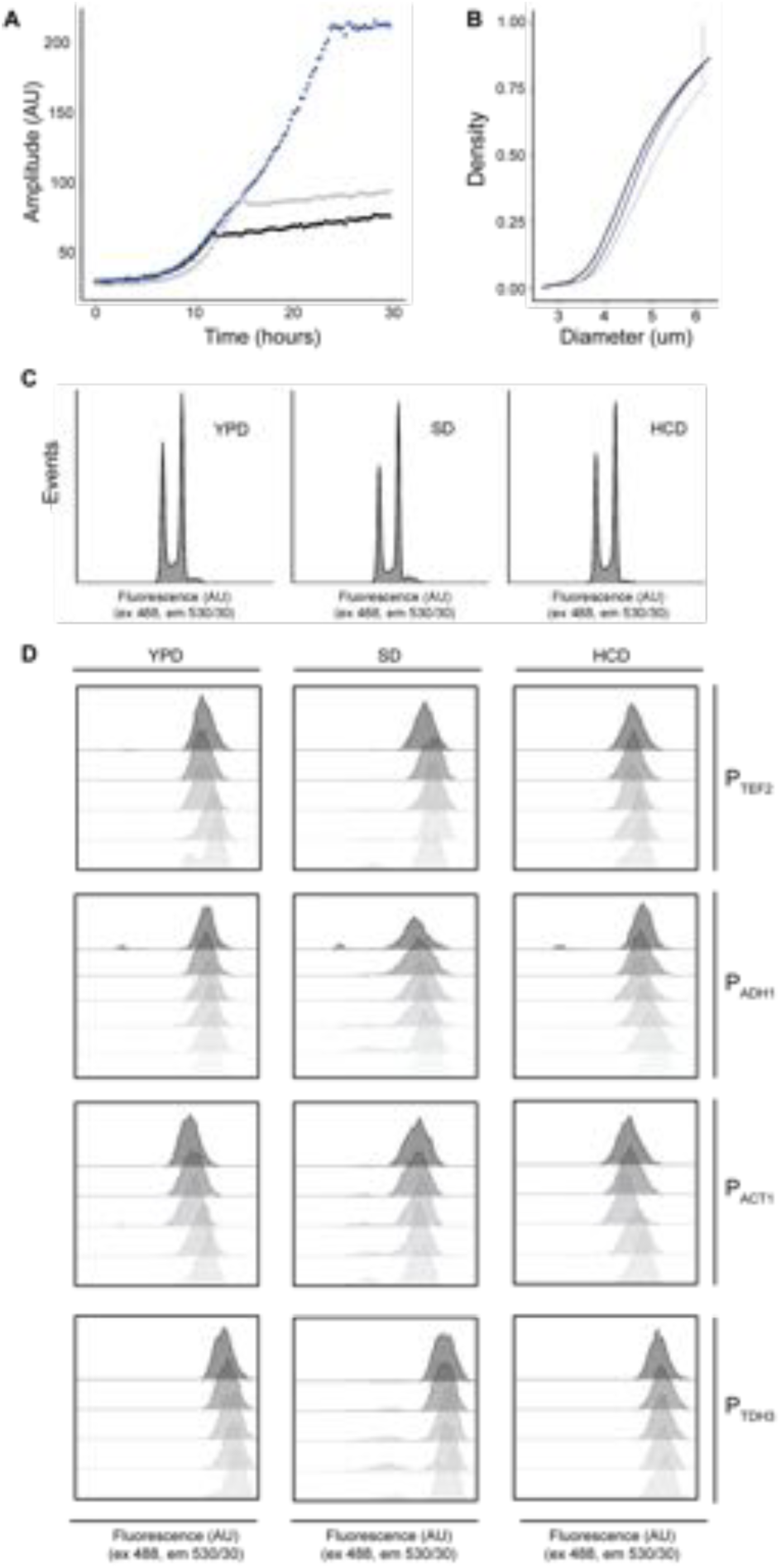
Characterisation of HCD media. (A) Comparison of the final optimised media (HCD; blue) with the initial media (black) and YPD (grey). (B) The cell size of BY4741 cells from logarithmicaly growing cultures was measured using a Coulter counter. The empirical cumulative distribution function of the measured cell size of SD (black), YPD (grey), and HCD (blue) is shown. (C,D) Cells from an exponentially growing culture were serially diluted 1:2 and seeded in a deep-well plate and incubated at 30°C at 300 rpm in the indicated medium. (C) Cells in mid-log phase were harvested, fixed and the DNA stained with Styox green. DNA content (which is proportional to fluorescence) is shown. (D) Cells expressing Citrine from the indicated promoters were harvested from different growth phases (early log to stationary; top to bottom) and fluorescence was examined.

To confirm that cells grown in HCD medium exhibit normal cell cycle behaviour we first looked at DNA content in logarithmically growing cultures by staining the DNA with Sytox Green (Figure 4C). We found that cultures grown in HCD medium exhibit similar ratios of cells in G1 and G2/M as SD and YPD, with a similar number of cells in S-phase as YPD. This indicates that cells grown in HCD behave similarly to cells grown in YPD.

We further evaluated the cellular physiology by examining the activity of metabolic or growth rate regulated promoters during different phases of growth. We used a set of well-characterised, constitutive promoters: P_*TEF*2_, P_*ADH*1_, P_*ACT*1_, or P_*TDH*3_, driving the expression of the fluorescent reporter, citrine (Ottoz et al., 2014) (Figure 4D). We scored the respective expression levels using flow cytometry and interpret increases in cell-to-cell variation as a measure for differences in cellular physiology. We found that for all four promoters, expression at different cell densities in HCD and YPD medium lead to a unimodal distribution, except for P_*TEF*2_ which shows a remarkable bimodal expression peak at the diauxic shift in YPD and to a lesser extent in HCD. In contrast, all promoters show the tendency of bimodal expression in SD medium.

It is important that a medium is not only specific for a single strain background, e.g. the background in which the optimisation was performed. Therefore, we repeated the growth curve analysis for a set of common lab strain backgrounds and assessed their behaviour in HCD in comparison to YPD and SD. We tested CEN.PK, W303-1B, and the wild isolate RM11 (Brem et al., 2002). For W303-1B and CEN.PK it was necessary to supplement HCD with 400 *μ*g/m*ℓ* adenine and/or 400 *μ*g/m*ℓ* tryptophan. Akin to the S288C derivative BY4741, all strains exhibited similar growth to YPD, with extended growth phase and higher ODs pre-diauxic shift (Figure 5A). W303-1B shows a premature kink in the growth rate which most likely could be alleviated by optimising the Ade or Trp dosage.

**Figure 5.**
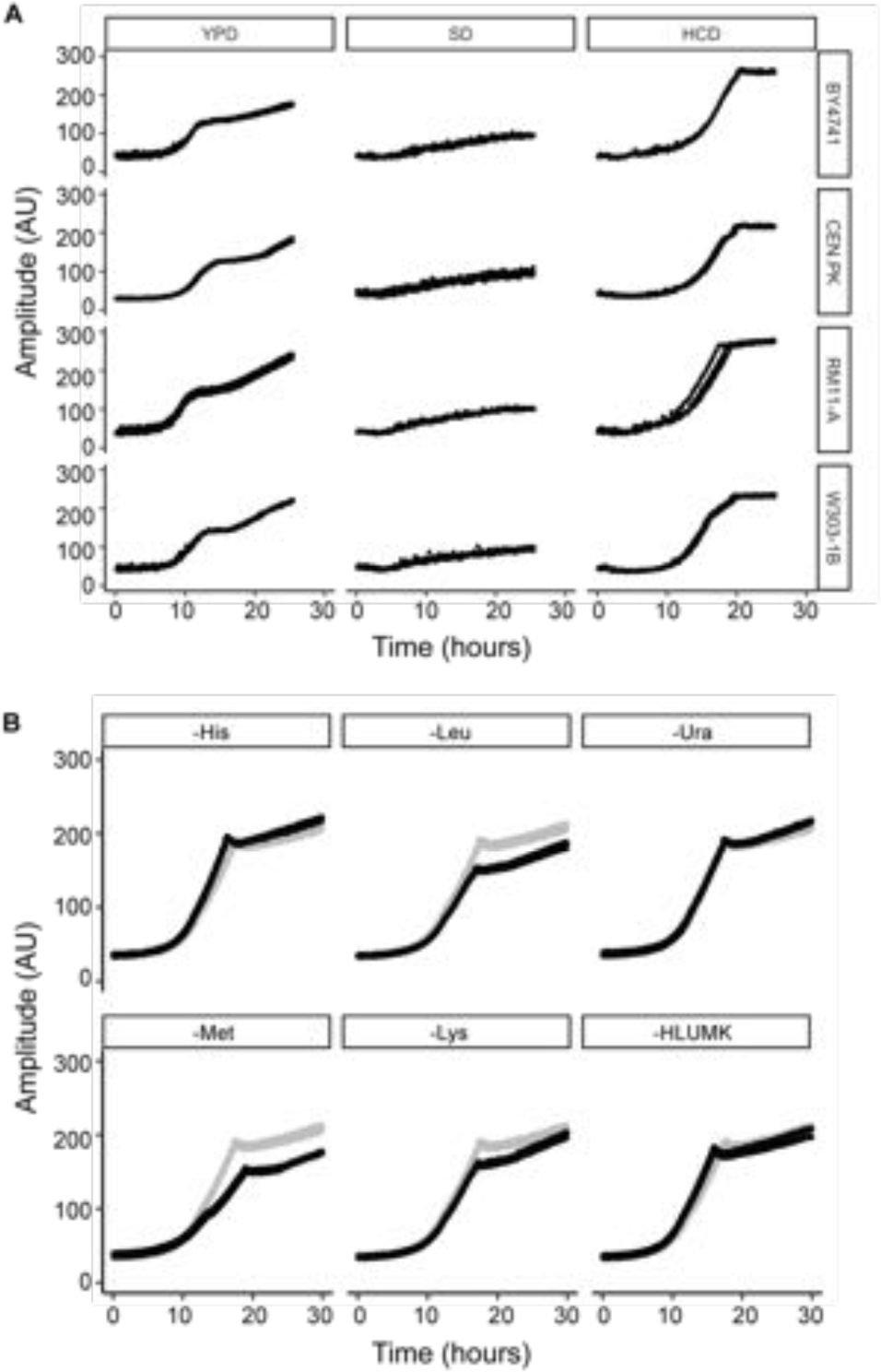
HCD is a versatile medium. 10 000 cells of the indicated strains (A) or 10 000 BY4741 cells (B) were seeded in a deep-well plate and incubated at 30°C at 300 rpm, in the indicated medium; growth in HCD media is shown in grey for comparison. Individual data points are plotted from at least 3 replicates.

We next wanted to confirm that HCD medium can be used with auxotrophic markers, which are essential tools in yeast research. They are used to maintain plasmids and to select for genomic integrations, and can vary among strain backgrounds. As such, we examined modified HCD media lacking specific amino acids and grew the prototrophic strain FY4 in HCD lacking one amino acid supplement at a time (denoted HCD-aa) or lacking amino acids corresponding to several common auxotrophic markers (denoted HCD-HLUMK) in the Biolector (Figure 5B). In general, the HCD dropout media showed only minimal deviation from the fully supplemented growth with the exception of medium lacking methionine (HCD-met), which grows slower, but still reaches a high cell density. HCD-Leu and -Lys showed a premature saturation, but identical growth rates throughout the growth curve.

### HCD medium allows for assay miniaturisation

One of the motivations for designing a defined high cell density medium was to facilitate the miniaturisation of assays such that assays classically performed in flasks could be performed in a more efficient manner using multi-well plates. As a proof of concept, we measured the secretion of *α*-amylase, a hard to secrete protein requiring the optimised media for detectable secretion (Wittrup and Benig, 1994; Tyo et al., 2012). *α*-Amylase is a relatively large protein, has an odd number of cysteines, and is glycosylated, all three characteristics disfavouring an efficient secretion. Expression of this protein therefore leads to a considerable amount of oxidative stress (Tyo et al., 2012).

We grew BY4741 cells harbouring a plasmid expressing *α*-amylase with a synthetic secretion signal (Tyo et al., 2012) in multi-well deep well plates in 600 *μℓ* of HCD or SD medium lacking uracil. We harvested at different growth phases and examined amylase activity in the supernatant by monitoring the release of p-nitrophenol (pNP) caused by amylase cleavage of ethylidene-pNP-G7 (Figure 6A). We were not able to detect any *α*-amylase activity at any stage of growth in SD medium. In contrast, we observed significant activity in the supernatants of cultures in stationary phase and late-log phase of cells grown in HCD medium. As expected, we observed more amylase secretion in growing cells compared to those in stationary phase.

**Figure 6.**
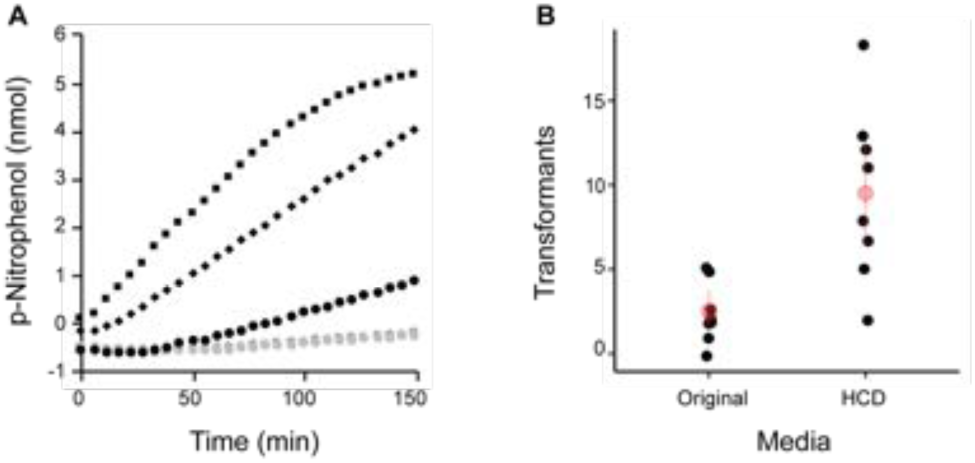
HCD facilitates miniaturisation of assays. (A) BY4741 cells expressing *α*-amylase from the plasmid (pYapAmyGPD) from an exponentially growing culture were serially diluted 1:2 and seeded in a deep-well plate and incubated at 30°C at 300 rpm in HCD-ura (black) or SD-ura (grey) overnight. The culture supernatant was sampled from different growth phases (early log (circles), late-log (squares), or stationary (diamonds) and *α*-amylase secretion was examined as function of the cleavage of ethylidene-pNP-G7 to yield p-Nitrophenol (OD_405_). (B) BY4741 cells containing a shuttle vector harbouring KANMX were grown in the indicated media with 200 *μ*g/m*ℓ* G418, plasmid was isolated and transformed into *E. coli*. The number of colonies obtained from seven independent plasmid isolations and subsequent transformations was determined (black points), the mean (red points) and standard deviation (red lines) are indicated.

We next compared the amount of plasmid that can be obtained from a yeast culture grown in the starting media and in HCD. To this end, we grew 1 m*ℓ* of a 1:25 fold diluted culture for 8h in a 96-well deep well plate in the initial starting media (i.e already optimised compared to SD) and HCD. We isolated the plasmid as described in material and methods and indirectly measured the amount of recovered plasmid by counting the colonies of *E. coli* cells transformed with the plasmid (Figure 6B). We found that the number of transformants correlated well with the optical densities of the input and took this as an indication that the optimised media was behaving as expected.

## Discussion

Miniaturising biochemical assays in *S. cerevisiae* is difficult as the commonly used strains, plasmids and auxotrophic markers only grow in a medium with a short exponential phase reaching small cell numbers at the diauxic shift. Using response surface methodology, we obtained an altered version of the commonly used synthetic medium with modified composition of its ingredients. We termed the medium HCD, for **h**igh **c**ell **d**ensity. Cultures grown in HCD medium reached at least three- and tenfold higher density at the diauxic shift compared to YPD and SD, respectively, and the balanced growth phase lasted up to 12h compared to 3–5h for the two standard media. HCD medium is compatible with widely distributed lab strains, as well as commonly used auxotrophic marker genes and combinations thereof. This is illustrated by the fact that the auxotrophic strain (BY4741) grows similarly in fully supplemented HCD as the prototrophic strain (FY4). The medium also supports positive selection as shown using the kanMX marker gene. The physiology of the cells growing in HCD medium seems normal, as their size remains similar to cells grown in standard conditions and the cell cycle distribution is similar to YPD. Importantly, the cell-to-cell variability of fluorescent proteins expressed from standard promoters is constant throughout the balanced growth phase. Additionally, the cell-to-cell variability is smaller than both SD and YPD grown cells, indicating that all cells experience a similar environment throughout the cultivation. Taken together, the results consistently showed that the HCD medium outperforms not only SD, but also YPD in terms of length of a balanced growth phase, cell numbers reached at the diauxic shift, and uniform protein expression, all while growing at a rate closer to YPD than SD.

Response surface methodology is widely used in process optimisation. Here, we used it to optimise the composition of a commonly used synthetic growth media to obtain a medium capable of supporting growth to high cell density. We performed two rounds of exploration and optimisation in the direction of steepest ascent. The method maximises a target value whose definition should reflect all desired properties. In our case, we realised that optimisation solely for cell density leads to slow growth and large cells. We therefore optimised for a combination of growth rate and cell density, while using an increase in cell size as an exclusion criteria. The method is therefore well suited for any process where the goal can be well quantified and written as a target value.

While the final medium composition is composed of “more of everything", the optimisation illustrated that the composition has to be well balanced otherwise unfavourable changes in cellular physiology will manifest. Compared to the standard SD, the final medium is made up of three-times more glucose, but roughly nine- and twelve-fold more MSG and amino acid pool, respectively. Therefore, to keep the cellular morphology and cell cycle distributions unchanged at higher densities, the addition of glucose needed to be overcompensated by the addition of amino acids as well as an increase of YNB and inositol. Interestingly, the first iteration of the optimisation pointed in this direction by lowering the amount of glucose and YNB while increasing the amount of amino acids and inositol provided. In the second iteration the pattern was similar but less pronounced, as we initially created an increase in biomass and cell size as well as a slow down of growth by increasing everything. Lowering the rate of glucose increase while increasing the amount of YNB, MSG and inositol corrected the optimisation to a higher cell density without increasing the cell size or distorting cellular physiology. HCD medium thus represents a finely tuned medium for balanced cell growth to high densities.

Importantly, cells grown as a small batch in HCD medium showed an increased secretion of amylase. The increase is larger than expected based on the increased cell density likely due to the more uniform growth of every member of the population as well as an increase of protein production per time as the cells grow faster in HCD than in SD. We therefore successfully miniaturised *α*-amylase secretion, a difficult to produce protein and hence hard to scale down. Additionally, we showed that purification of plasmid DNA is feasible using a standard 96 glass filter well plate. By extension, we assume that HCD will allow for efficient protein assays obtained from small volumes grown in 96 well plates. As the different HCD media are based on the common ingredients found in every yeast lab, we expect them to be of instantaneous use when large cell numbers are required, but cultivation is only feasible in small vessels.

## Acknowledgements

The authors thank Grant Brown for carefully reading and commenting on the manuscript. This work was funded by the ETH (TR, HMK, FR) and the NCCR Molecular Systems Engineering (FR). We thank Yannick Schmid for initial work, and Sven Panke and Jörg Stelling for helpful discussions and support.

## Material and Methods

### Yeast strains

Different strains were used and Table 3 gives an overview of the source and the relevant genomic modification.

### Standard media recipes

Standard yeast media (YPD and SD) compositions were used as described in Sherman (1991).

### Stock Solutions

The individual chemicals for the medium optimisation were provided as concentrated stock solutions. The different pools of chemicals and the individual concentrations are given in Table 4. *All components were filter-sterilised

### Chemicals

All chemicals were obtained from Sigma-Aldrich (Buchs, Switzerland) unless otherwise indicated.

### Growth experiments

Strains were grown overnight at 30°C in 2 m*ℓ* of the indicated medium. The next day 10 000 or 20 000 cells were seeded in 800 *μℓ* of the same medium in a 48-well FlowerPlate (without optodes) (m2p-labs, Bäsweiler, Germany). The plate was covered with a gas-permeable sealing film (m2p-labs) and placed in a BioLector device (m2p labs) incubated at 30°C, with 20.95% O_2_, ≥ 85% humidity, shaken at 1 300 rpm and biomass was measured as a function of back scatter at 600 nm every 15 minutes.

### Measurement of cell size and number

100 *μℓ* of cell suspension was added to 10 m*ℓ* of PBS and cell size distribution and cell number was measure in a Z2 Coulter Counter (Beckman Coulter, Nyon, Switzerland), with a 100*μ*m orifice. Very dense cultures cells were additionally diluted 1:10 in PBS.

### Analysis of cell-cycle behaviour

Strains were grown overnight at 30°C in 2 m*ℓ* of the indicated medium. The next day the culture was diluted 1:20 in the same medium and grown for 6-7 hours at 30°C. 600 *μℓ* of culture was serially diluted (1:2) across 8 or 12 wells of a 96-well deep well plate (Kuhner, Birsfelden, Switzerland). The plate was covered with a gas-permeable lid and placed in an ISF1-X (50 cm diameter) Kuhner shaker and incubated at 30°C shaken at 330 rpm overnight. The next day 100 *μℓ* of cells were transferred to a 96-well MTP (Greiner Bio One, Kremsmünster, Austria) diluted with PBS and fluorescence of strains was analysed using a Fortessa flow analyser (Becton Dickinson, Allschwil Switzerland). For DNA staining cells were fixed by resuspending in 500 *μℓ* of cold 70% ethanol and incubated 15 minutes at RT. Cells were harvested and resuspended in 500 *μℓ* of 2 mg/m*ℓ* Proteinase K in 50 mM Tris pH 7.5 and incubated for 50 minutes at 50°C. Cells were harvested and resuspended in 500 *μℓ* 0.2 mg/m*ℓ* RNaseA in 50 mM Tris pH 7.5 for 4 hours at 37°C. Cells were harvested and resuspended in 500 *μℓ* FACS buffer (200 mM NaCl, 78 mM MgCl_2_, 200 mM Tris pH 7.5) and stored at 4°C. Before analysis 100 *μℓ* of cells were added to 1 m*ℓ* of 1 *μ*M Sytox Green (Thermo Fisher, Basel, Switzerland) in 50 mM Tris pH 7.5.

### Amylase secretion assay

BY4741 harbouring pYapAmyGPD (Tyo et al., 2012) expressing *α*-amylase were grown overnight at 30°C in 2 m*ℓ* of the media lacking uracil. The next day the culture was diluted 1:20 in the same medium and grown for 6-7 hours at 30°C. 600 *μℓ* of culture was serially diluted (1:2) across 12 wells of a 96-well deep well plate (Kuhner, Birsfelden, Switzerland). The plate was covered with a gas-permeable lid and placed in an ISF1-X (50 cm diameter) Kuhner shaker and incubated at 30°C shaken at 330 rpm overnight. The next day cells were pelleted for 3 min, 2 000 × *g* in an Eppendorf 5920R centrifuge (Vaudaux-Eppendorf AG, Schönenbuch, Switzerland). 50 *μℓ* of supernatant was transferred to a 96-well multi-well plate (Greiner Bio One, Kremsmünster, Austria). The amylase assay was performed using the Amylase Activity Assay Kit from Sigma (MAK009), all steps were performed according the manufacturer’s instructions. OD405 was measured using a Tecan Infinite M Nano Absorbance Reader (Tecan Group Ltd., Männedorf, Switzerland).

### Plasmid rescue and transformation

Overnight cultures of BY4741 harbouring FRP2061 (a 13kb plasmid with a dual promoter Kanamycin cassette for resistance in both *E. coli* (kanamycin) and *S. cerevisiae* (G418)) were diluted 1:25 in 1 m*ℓ* in deep-well plates (Kuhner, Birsfelden, Switzerland). The plate was covered with a gas-permeable lid and placed in an ISF1-X (50 cm diameter) Kuhner shaker and incubated at 30°C shaken at 330 rpm for 8 hours in the indicated media with 340 *μ*g/m*ℓ* G418. Samples were harvested and stored at 4°C in 50 *μℓ* spheroplasting solution (per 1 m*ℓ* 1M sorbitiol, 6 *μℓ* 1 M DTT and 15 *μℓ* 10 mg/m*ℓ* 100T zymolyase) overnight. The next day the samples were incubated 2 hours at 37°C and spheroplasting was verified microscopically. 50 *μℓ* Qiagen P1 buffer was added, followed by 50 *μℓ* Qiagen P2 buffer and 70 *μℓ* Qiagen N3 buffer (Qiagen, Hilden, Germany). The sample was transferred to 96-well PVDF plate, stacked on top of a 96-well glass filter plate, on top of 96-well collection plate and spun 3 000 × *g*, 10 minutes in an Eppendorf 5920R centrifuge (Vaudaux-Eppendorf AG, Schönenbuch, Switzerland). The glass filter plate was washed with cold 70% EtOH and spun at 3 000 × *g* for 3 minutes and washed with cold 100% EtOH and spun at 3 000 × *g* for 3 minutes. Plasmid was eluted with 25 *μℓ* TE buffer (pH = 8) into clean 96-well plate by spinning at 3 000 × *g* for 3 minutes. 5 *μℓ* of the eluate was transformed into 45 *μℓ* chemically competent *E. coli* DH5*α* cells and plated on LB + 50 *μ*g/mℓ.

## Supplementary Text - Response surface methodology

We give a brief overview over the response surface techniques. We denote by *y* the measured response, which for our purposes is either the increase in optical density (OD): *y* = ΔOD or the increase in OD and duration: *y* = ΔOD − 2 · Δ*t*. We describe the response surface by the expected response resulting from setting the concentrations of five medium components: glucose concentration (with concentration *c*_1_), the concentration or amounts of the two nitrogen sources, (*c*_2_, *c*_3_), YNB (*c*_4_), and inositol (*c*_5_). The response surface is then a function *y* = *f* (*c*_1_, *c*_2_, *c*_3_, *c*_4_, *c*_5_), but its form *f* () is unknown. The response surface methodology is a systematic approach for designing experiments to iteratively approach the maximum of this function.

The response surface methodology was first developed to determine optimal conditions in chemical experiments (Box and Wilson, 1951), and then adapted for a variety of other applications. A very readable classical account is given in (Box et al., 2005). We used the package rsm (Lenth, 2009) for the statistical software R (R Core Team, 2019) for all experimental designs and calculations, but many other statistical software packages also provide corresponding functionality.

We consider a specific medium composition 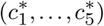 as our reference point from which we start the optimisation. The goal is to first explore the response surface around this reference point to determine its local shape, and then to calculate the direction in which the surface increases fastest, known as the canonical path (if it is curved) or the gradient of steepest ascent (if it is a straight line).

### Central composite designs for exploration

In order to explore and determine the local shape of the response surface, we specify, separately for each explanatory variable *c*_*i*_, a value Δ_*i*_ and define two new values: one lower 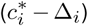, the other higher 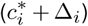 than the reference value 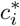. The maximal changes Δ_*i*_ thus determine the vicinity around the reference point. If the chosen vicinity is too small, many iterations of RSM will be required, and inaccuracies can occur if the difference in response values at the low and high points are small compared to the measurement noise. On the other hand, choosing a too large Δ_*i*_ can result in over-approximation of the response surface, and optimal values might be missed. The definition of the vicinity around the reference point therefore requires attention. In practice, logarithmic changes are often a good starting point, using one-half and double the amount of each component. For our first iteration, the low and high values are shown together with the reference point in the first three rows of Table 1.

We then determined three sets of new medium compositions at which we experimentally determined the response value *y*: (i) the central point, (ii) the factorial points, and (iii) the axial points. Together, these form a *central composite design (CCD)*, illustrated for three components in Figure 1.

The central point is simply the reference point, and we take measurements for several replicates at this point. These replicates serve three functions: first, they provide a direct check of the reproducibility of the reference condition, second, they allow us to estimate the standard deviation of the measurements directly, and third, they allow us to separate lack-of-fit of the statistical model from measurement error and determine how good the local approximation of the response surface is.

The factorial points are all combinations of low and high values for the five components; statistically, they correspond to a full factorial design. They form the corners of a five-dimensional box parallel to the coordinate axis, with the central points in the centre of this box. Importantly, they will allow estimating *interactions* between components, for example, if an increase in the first component only yields an increase in the response if the second component is decreased simultaneously.

Finally, we define two axial points for each component by using its low and high value while keeping all other components at their reference value. A line through these two points intersects the box formed by the factorial points in the centre of the corresponding box’s facets.

The choice of a reference condition and the central composite design is illustrated in Figure 1B-i for two components. For computational reasons and to aid interpretation, RSM uses a new coordinate system (*x*_1_,…, *x*_5_) in which we perform the estimation and calculations: we shift the origin to the coordinates of the reference point, and we scale each axis such that the low and high values for that component coincide with the values ±1. Specifically, the new value *x*_*i*_ for the *i*th component is

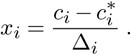

This new coordinate system is shown in Figure 1B-ii; note that the previous CCD ‘box’ is now a square and that the coordinate origin is at the starting condition. In these new coordinates, each centre point has coordinate (0,…, 0), each factorial point has coordinate (±1,*…, ±*1), and the axis points for the third component, for example, have coordinates (0, 0, ± 1, 0, 0). Thus, the factorial and axial points lie on the corners of a five-dimensional unit cube, and in the centre of each of its facets, respectively.

In the general case of *k* explanatory variables, we require 2^*k*^ factorial points and 2 × *k* axial points, and we are free to choose the number of replicates *n* of the centre points. We chose *n* =6 centre points to provide sufficient replication, and thus have a full design with 32 + 10 + 6 = 48 experimental conditions. This number can be reduced substantially by employing ideas from factorial designs and only consider every second corner of the unit cube (a half-fraction of the full factorial design), resulting in 16 + 10 + 6 = 32 experimental conditions.

### Model

Most commonly, a second-order model is used to locally approximate the response surface in the vicinity of a reference point. For five components, this model is given by

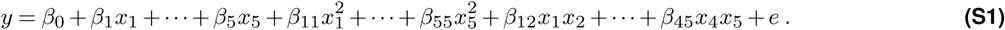

In the scaled and shifted coordinate system, the parameter *β*_0_ is thus the expected response at the reference point. The parameters *β*_*i*_ describe a linear first-order model that corresponds to a hyperplane. The purely-quadratic parameters *β*_*ii*_ allow independent curvature along each axis, and the two-way-interaction-parameters *β*_*ij*_ allow the effect on the response *y* of one component *i* to depend on the value of a second component *j*. Finally, the residual error *e* is assumed to be normally distributed with mean zero and constant variance.

The parameters can be estimated from the data of a CCD by least-squares, for example. Especially in the first iterations of the experiments, the approximated response surface might have little global resemblance with the true surface but should be accurate enough in the vicinity of the reference point to decide on the next experimental conditions to test. This is illustrated in Figure 1B-iii, where the very local CCD leads to an approximated response surface that is almost a plane that nevertheless describes the local properties of the true surface reasonably well.

### Pursuing the canonical path of steepest ascent

Once the parameters of the model are estimated, we calculate the path along the response surface that give the steepest increase in response value. In the simplest case, this is a straight line, known as the gradient of steepest ascent. For second-order models such as (S1), the steepness will change along the surface due to the curvature, and we instead calculate the *canonical path*, which is a curved line along which we achieve the steepest increase in response value.

The next experimental conditions are then chosen along the canonical path, at regular distances from the reference point. Here, we used distances 0.0, 0.5, 1.0, up to 5.0; depending on the direction, points at distances of 1 to about 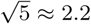 or higher are outside the region initially explored by the CCD and the predicted response values are then based on extrapolation. Especially in the first iteration(s), the response surface approximation can be relatively inaccurate, and with increasing distance, the measured values along the canonical path might then start deviating substantially from those predicted for the corresponding medium composition. We observe this phenomenon in the first iteration of our illustration (Figure 1B-iii), where the canonical path first increases and then decreases again, yielding an intermediate optimum at the star-shaped point. The same phenomenon can be seen in our actual data (Fig. 2C).

### Sequential experimentation

The two steps of locally exploring the response surface and building an approximate model, followed by determining the canonical path that yields the fastest increase in response and sampling the surface experimentally along this path can now be iterated several times. The result of a second iteration is shown in Figure 1B-iv, where the intermediate optimum found along the canonical path of the first iteration is taken as the new center point, a CCD is designed around this point, and the measurements taken at the corresponding experimental conditions. This yields a new (local) approximation of the response surface, for which a new canonical path is calculated and pursued. In practice, the increase in response often starts to slow substantially after very few iterations of the two steps, and 2–4 iterations are fairly typical until the experimental effort does no longer justify the anticipated gain.

## Supplementary figures & tables

**Table S1.**
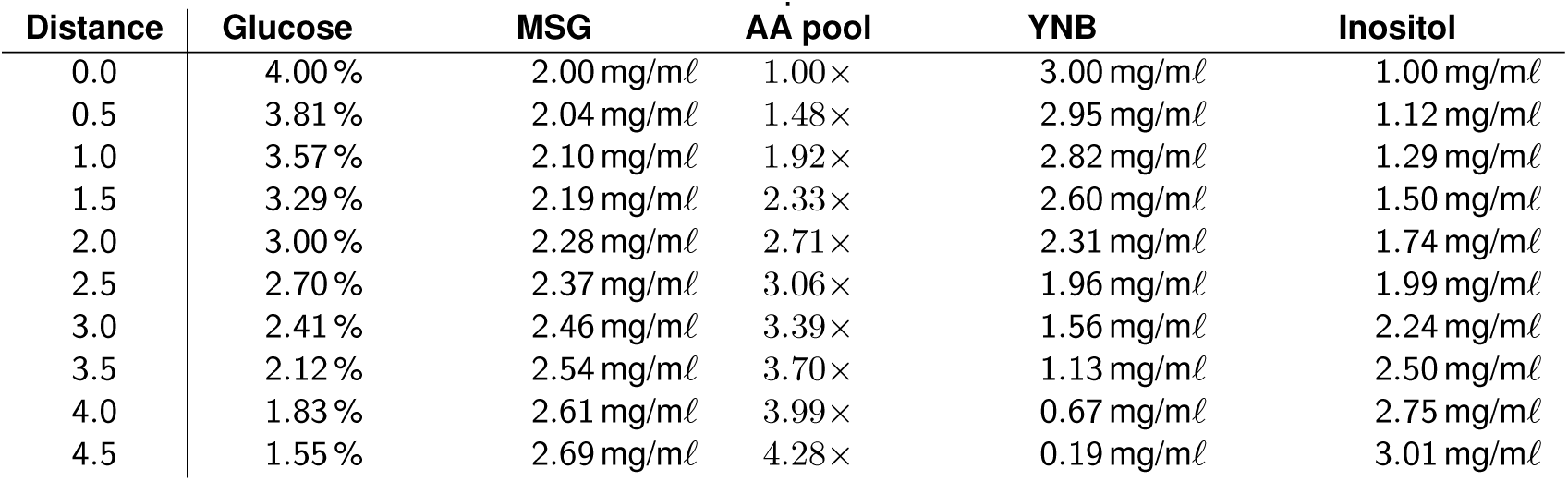
Media compositions along the first steepest ascent.

| Distance | Glucose | MSG | AA pool | YNB | Inositol |
| --- | --- | --- | --- | --- | --- |
| 0.0 | 4.00 % | 2.00 mg/ml | 1.00× | 3.00 mg/ml | 1.00 mg/ml |
| 0.5 | 3.81 % | 2.04 mg/ml | 1.48× | 2.95 mg/ml | 1.12 mg/ml |
| 1.0 | 3.57 % | 2.10 mg/ml | 1.92× | 2.82 mg/ml | 1.29 mg/ml |
| 1.5 | 3.29 % | 2.19 mg/ml | 2.33× | 2.60 mg/ml | 1.50 mg/ml |
| 2.0 | 3.00 % | 2.28 mg/ml | 2.71× | 2.31 mg/ml | 1.74 mg/ml |
| 2.5 | 2.70 % | 2.37 mg/ml | 3.06× | 1.96 mg/ml | 1.99 mg/ml |
| 3.0 | 2.41 % | 2.46 mg/ml | 3.39× | 1.56 mg/ml | 2.24 mg/ml |
| 3.5 | 2.12 % | 2.54 mg/ml | 3.70× | 1.13 mg/ml | 2.50 mg/ml |
| 4.0 | 1.83 % | 2.61 mg/ml | 3.99× | 0.67 mg/ml | 2.75 mg/ml |
| 4.5 | 1.55 % | 2.69 mg/ml | 4.28× | 0.19 mg/ml | 3.01 mg/ml |

**Table S2.**
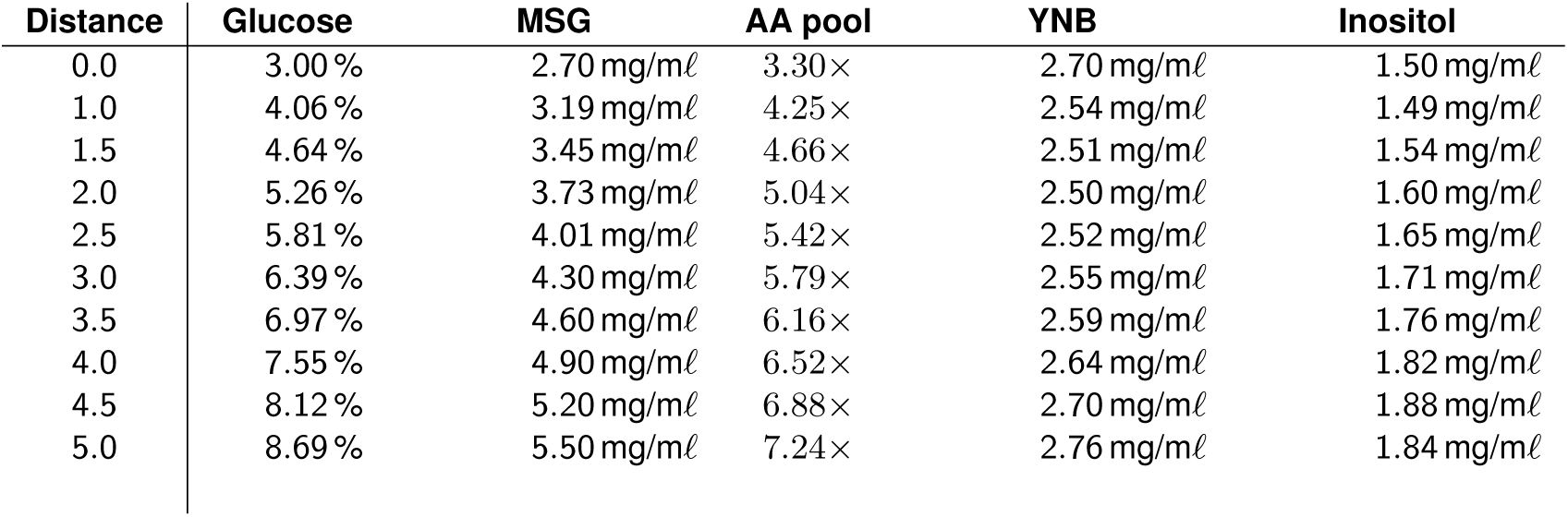
Media compositions along the second steepest ascent. Optimisation based on OD only.

| Distance | Glucose | MSG | AA pool | YNB | Inositol |
| --- | --- | --- | --- | --- | --- |
| 0.0 | 3.00 % | 2.70 mg/ml | 3.30× | 2.70 mg/ml | 1.50 mg/ml |
| 1.0 | 4.06 % | 3.19 mg/ml | 4.25× | 2.54 mg/ml | 1.49 mg/ml |
| 1.5 | 4.64 % | 3.45 mg/ml | 4.66× | 2.51 mg/ml | 1.54 mg/ml |
| 2.0 | 5.26 % | 3.73 mg/ml | 5.04× | 2.50 mg/ml | 1.60 mg/ml |
| 2.5 | 5.81 % | 4.01 mg/ml | 5.42× | 2.52 mg/ml | 1.65 mg/ml |
| 3.0 | 6.39 % | 4.30 mg/ml | 5.79× | 2.55 mg/ml | 1.71 mg/ml |
| 3.5 | 6.97 % | 4.60 mg/ml | 6.16× | 2.59 mg/ml | 1.76 mg/ml |
| 4.0 | 7.55 % | 4.90 mg/ml | 6.52× | 2.64 mg/ml | 1.82 mg/ml |
| 4.5 | 8.12 % | 5.20 mg/ml | 6.88× | 2.70 mg/ml | 1.88 mg/ml |
| 5.0 | 8.69 % | 5.50 mg/ml | 7.24× | 2.76 mg/ml | 1.84 mg/ml |

**Table S3.** Media composition of the second steepest ascent. Optimisation based on OD and time

| Distance | Glucose | MSG | AA pool | YNB | Inositol |
| --- | --- | --- | --- | --- | --- |
| 0.0 | 3.00 % | 2.70 mg/ml | 3.30× | 2.70 mg/ml | 1.50 mg/ml |
| 0.5 | 3.48 % | 2.95 mg/ml | 3.30× | 2.65 mg/ml | 1.45 mg/ml |
| 1.0 | 4.02 % | 3.24 mg/ml | 4.25× | 2.65 mg/ml | 1.51 mg/ml |
| 1.5 | 4.55 % | 3.56 mg/ml | 4.66× | 2.69 mg/ml | 1.57 mg/ml |
| 2.0 | 5.08 % | 3.90 mg/ml | 5.04× | 2.77 mg/ml | 1.64 mg/ml |
| 2.5 | 5.60 % | 4.25 mg/ml | 5.42× | 2.87 mg/ml | 1.71 mg/ml |
| 3.0 | 6.12 % | 4.62 mg/ml | 5.79× | 2.99 mg/ml | 1.78 mg/ml |
| 3.5 | 6.63 % | 4.98 mg/ml | 6.16× | 3.12 mg/ml | 1.86 mg/ml |
| 4.0 | 7.13 % | 5.36 mg/ml | 6.52× | 3.26 mg/ml | 1.93 mg/ml |
| 4.5 | 7.62 % | 5.74 mg/ml | 6.88× | 3.42 mg/ml | 2.01 mg/ml |
| 5.0 | 8.12 % | 6.12 mg/ml | 7.24× | 3.57 mg/ml | 2.09 mg/ml |

**Figure S1.**
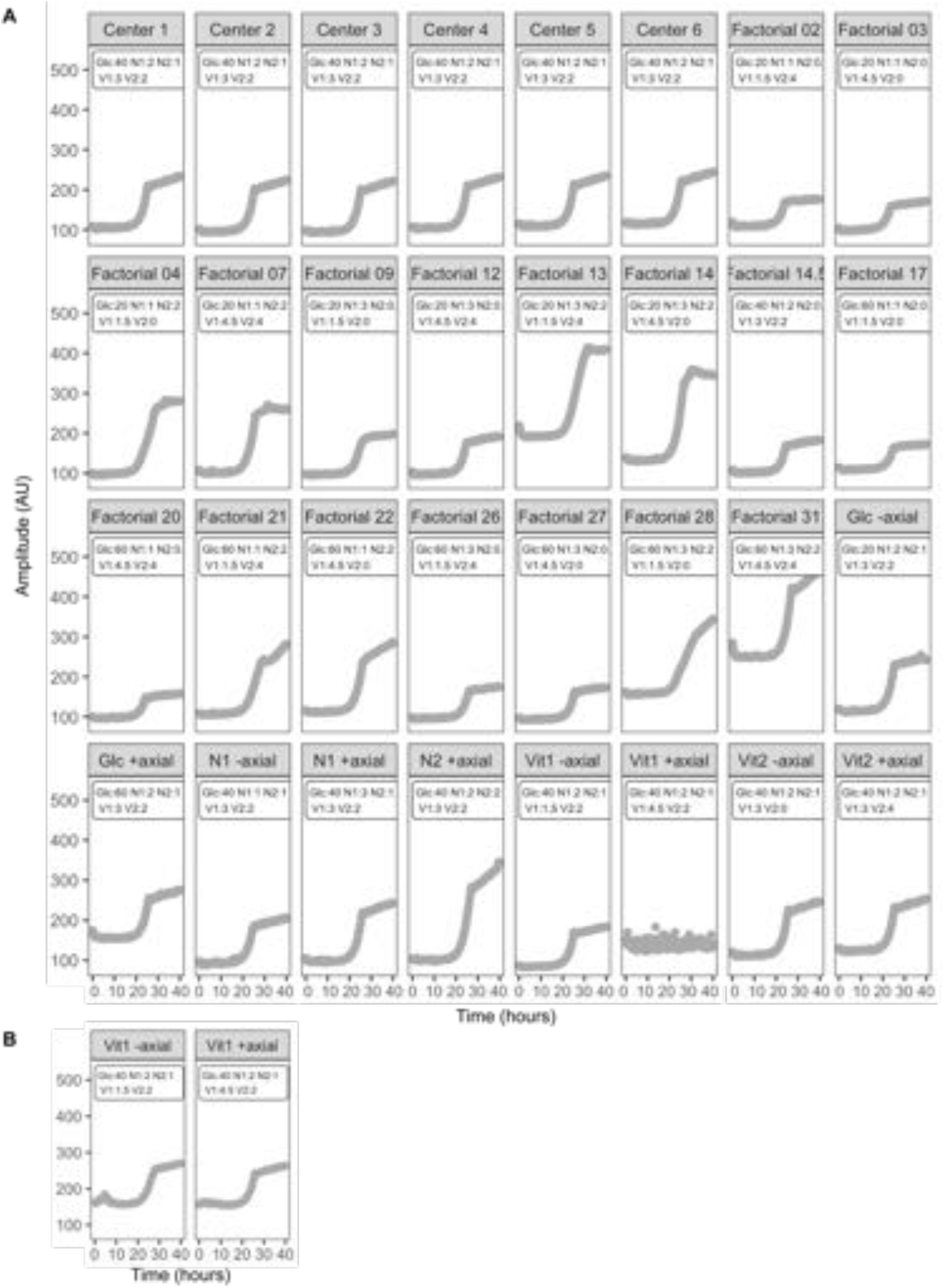
First exploration of media conditions. (A) Growth curves of BY4741 cells grown in the indicated media compositions. (B) Curves obtained for Vit1 − and + axial in an independent experiment. Numbers indicate volume in *μℓ* added of the stock solutions of each component in 1 m*ℓ* of media.

**Figure S2.**
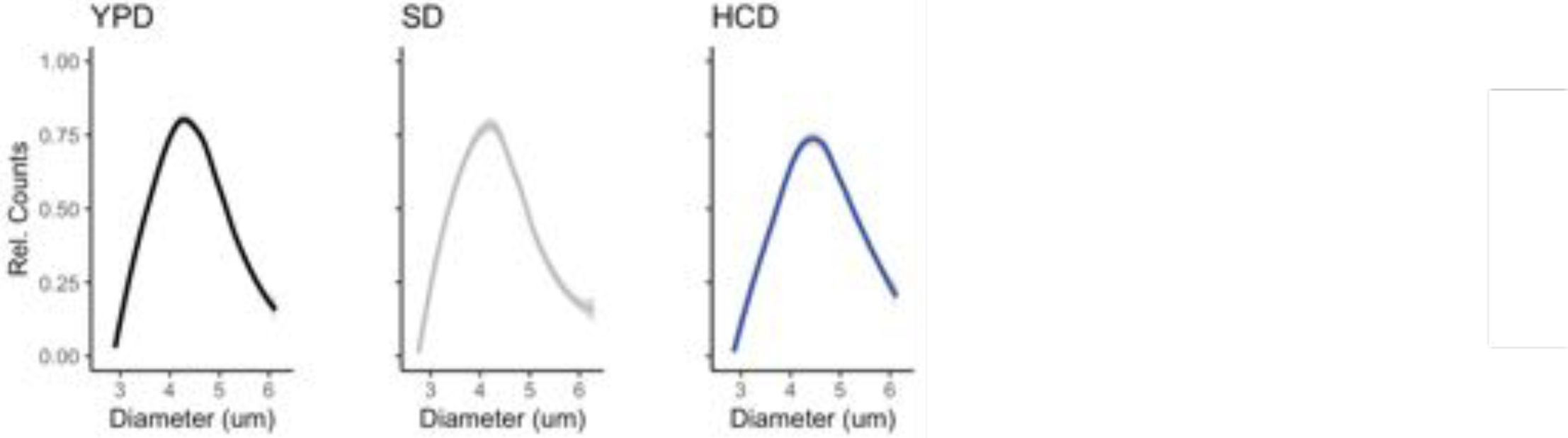
Cell size distribution. The size of BY4741 cells in mid-log phase grown in the indicated media were measured using a Coulter counter.

